# Analysis of Bone Healing with a Novel Bone Wax Substitute Compared to Bone Wax in a Porcine Bone Defect Model

**DOI:** 10.1101/236430

**Authors:** Tristan Tham, Keith Roberts, John Shanahan, John Burban, Peter Costantino

**Author notes:** **Lenox Hill Hospital (Black Hall Building)**, 130 East 77th Street, 10th Floor, New York, NY 10075. **Senior author:** Peter Costantino, MD –. **Co-authors:** Keith Roberts –, John Shanahan –, John Burban –. **Corresponding author:** Tristan Tham, MD –.

## Abstract

**Background:** Bone wax is used in surgery as a hemostatic device for bone. Despite its good functional capacity as a bone hemostat, Bone wax materials often have very poor long-term interactions with bone. This study describes a novel composite of hydroxyapatite (HA) and biodegradable poly-lactic acid (PLA) with wax-like handling properties (**OsteoStat**). The goal was to compare qualitative and quantitative measures between OsteoStat versus Bone wax.

**Methods:** The porcine critical size defect model was chosen in this study. OsteoStat and Bone wax were introduced into separate critical size defects located in the femur and humerus of a single porcine specimen. After a duration of 6 weeks, the defect sites were harvested for clinical, histological, and histomorphometric analysis.

**Results:** Both groups had effective hemostatic action when introduced into the defects. Analysis of the histomorphometric data revealed that the amount of new bone was significantly greater at 6 weeks in the OsteoStat group (38.05%) versus the Bone wax group (11.88%), p=0.028. OsteoStat also demonstrated less soft tissue and less test material remaining in the defect sites; however, this was not statistically significant.

**Conclusions:** We speculate that the incomplete biodegradation of Bone wax as well as its intrinsic inflammatory properties may have retarded osseous regeneration and promoted fibrosis. In contrast, well known biodegradation pathways for PLA combined with the HA component of OsteoStat may have accounted for the positive results of OsteoStat compared to Bone wax. It is important that bone hemostat substances have biocompatible, osteoconductive, hemostatic, as well as good handling properties.

## Background

Bone wax, first described by Sir Victor Horsley as an antiseptic in 1892, is now commonly used in surgery as a hemostatic device.^1,2^ In cardiothoracic surgery, the main indication for Bone wax is extensive bleeding from the sternal bone marrow. During sternotomy procedures, intraoperative exposure of the cut surfaces of the sternum, together with concurrent anticoagulant therapies increases the patients’ risk of bleeding into the sternal wound. Surgeons try to minimize such bleeding as the eventual intrasternal hematoma collection could serve as a nidus for bacterial infection, an uncommon but serious sequalae of cardiothoracic surgery.^3^ In order to prevent extensive hematoma formation, surgeons apply topical hemostatic agents, such as Bone wax, to the cut sternal marrow.^4^ Surgical Bone wax consists of a naturally occurring substance, sterilized *Cera Alba* (honeybees wax), and is usually mixed with paraffin wax which acts as a softening agent to enhance handling characteristics.

Since Bone wax is comprised of paraffin wax and esterified fatty acids, it is highly hydrophobic. This enables Bone wax to serve as a physical barrier against blood, occluding the bleeding channels and achieving hemostasis by a tamponade and blood stasis effect. However, this hydrophobic property in conjunction with limited enzymatic degradation of waxes in the human body prevents appreciable rates of absorption and/or excretion post-surgical application.. It is well documented that Bone wax impairs optimal bone formation and healing of sternotomies, ^3,5^ which could be due to, in part, physical inhibition of osteoblast and osteocyte migration to the site of bony injury. Furthermore, the nature of Bone wax which makes it highly resistant to degradation has also been linked with infection,^6,7^ although large randomized studies have found the infection link to be inconclusive.^8^ Since intraoperative bone bleeding can be heavy, physicians must weigh the benefits of bone hemostasis using Bone wax versus the risk of decreased bone healing and other complications such as infection.

It is important that Bone wax like substances have biocompatible, osteoconductive, hemostatic, as well as good handling properties. Several alternative materials have been reported in the literature such as PEG/collagen,^9^ polyorthoester,^10^ fibrin-collagen,^11^ chitin-based material,^12^, and gelfoam.^13^ However, none of these alternative materials have yet seen widespread adoption, suggesting that a material which meets the effective hemostatic qualities of Bone wax together with good osseous integration and affordability has not been met.

Herein we describe OsteoStat (Hemostasis LLC, MN, USA), a novel composite of hydroxyapatite (HA) and biodegradable poly-lactic acid (PLA) with wax-like handling properties. HA (*Ca10(PO4)6(OH)2*) is a biomaterial similar to the mineral component of natural bone, and exhibits good osteoconductivity.^14^ Similar to Bone wax, the OsteoStat creates a physical barrier which is its primary mechanism of hemostasis. Unlike Bone wax, we hypothesized that the OsteoStat would, due to its HA content, have an enhanced bone healing profile in addition to hemostatic qualities. The goal of this experiment was to compare the widely used cardiothoracic surgery hemostatic agent, Bone wax, versus OsteoStat. We investigated qualitative and quantitative measures of bone healing between the two materials.

## Materials & Methods

Details of animal husbandry, diet, care, monitoring, health, and well-being as well as measures to alleviate suffering were all performed in accordance with International Organization for Standardization document ISO 10993-11:2006: Biological evaluation of medical devices.

This experiment was conducted in a Sichuan Province Experimental Animal Management Committee Accredited Facility in China. Ethics approval for animal experimentation was approved by the animal ethics board of the Sichuan Province Experimental Animal Management Committee of China.

A porcine bone defect model was used for this study. One female young adult Landrace pig weighing 52.8 kg was used for this study, with an acclimation period of 7 days. Two groups of holes were drilled into the femur and humerus and filled with either OsteoStat or Bone wax. After a period of 6 weeks, qualitative (clinical and histology) and quantitative (histomorphometry) analyses were performed on the drilled sites.

### Surgical Procedure

The animal was prepared for operation under general anesthesia. Intraperitoneal pentobarbital sodium was administered and the field of operation was then sterilized and selected at the right humerus and contralateral left femur. Tissue dissection was performed, exposing the underlying periosteum at these respective sites. Four test sites were chosen for the bone defect, 2 holes located in the right diaphyseal humerus and 2 holes located in the left diaphyseal femur. All test sites were drilled to have standardized intraosseous defects with a circumference of 3cm and equal depth. Care was taken to drill the cortical bone down to a similar depth in all the drill sites. One defect on each bone (humerus or femur) was filled with OsteoStat (Hemostasis LLC, MN, USA) and the other defect was filled with Bone wax W31G (Ethicon, NJ, USA). The surgical sites were closed in multiple layers.

The animal was given a healing period of 6 weeks before it was sacrificed. During the healing period, the animal displayed no signs of local infection and surgical site incisions were well healed. The animal was fully ambulant during the entire period and was effectively load bearing on all four limbs. After the healing period, the animal was sacrificed with an overdose of pentobarbital sodium. The right humerus and left femur diaphysis were cut to harvest the test sites as discrete blocks. On harvesting of the test sites, no signs of gross inflammation or necrosis were observed in any of the sites.

### Histology Preparation

The harvested blocks of tissue containing the test sites were fixed with a 10% neutral buffered solution of formalin for a period of 7 days. The blocks were then decalcified by a solution of mixed acid decalcification agent for 6 weeks. Next, the sections were dehydrated with ethanol gradient and prepared with paraffin. Hematoxylin and Eosin (H&E) stain was applied to the sections for histological observation at x40, x100, and x 400 magnification. Two sections were prepared from each of the 4 drilled sites, for a total of 8 sections for histological/histomorphometric analysis (4 sections for the OsteoStat defect and 4 sections for the Bone wax defect).

### Histomorphometric Evaluation

Two sections per drilled site were used for histomorphometric analysis, for a total of 4 sections per test material. Analysis was done at the x40 power magnification level, as this would give a balance between tissue resolution and reduction in section variability, which would be greatly biased at higher power magnification. Histomorphometric data was obtained by analyzing the magnified cross sections of the defect sites using National Institute of Health program software, ImageJ 1.48v. Areas of interest were measured in terms of pixel counts. Parameters delineated in the histomorphometric evaluation were % bone area, % soft tissue area, % test material, all of which combined would add up to approximately 100% of the defect size.

### Statistical Analysis

Histomorphometric data was analyzed by comparing the mean between the two test materials using Student’s t-test. Results were considered significant with a p value of less than 0.05.

## Results

### Clinical Evaluation

Upon drilling defects into the femoral and humeral diaphysis, bone bleeds were observed in all the bone defects. There was effective hemostasis in both materials. The animal remained healthy for the duration of the study with no post-operative or surgical site complications. The rest of the 6 week healing period was uneventful.

### Histology

A selection of images which are representative of the qualitative findings are included below.

#### X40power magnification

In the OsteoStat sites, new bone formation was observed, in addition to the soft tissue stroma. Furthermore, new vessel formation was observed in some of the sections **(Fig. 1).** In the Bone wax sites, fibrous soft tissue stroma was the predominating component. Some new bone can also be seen, and there is also the presence of undegraded test material. The amount of new bone appears visually less than in the OsteoStat sites **(Fig. 2).**

**Fig. 1.**
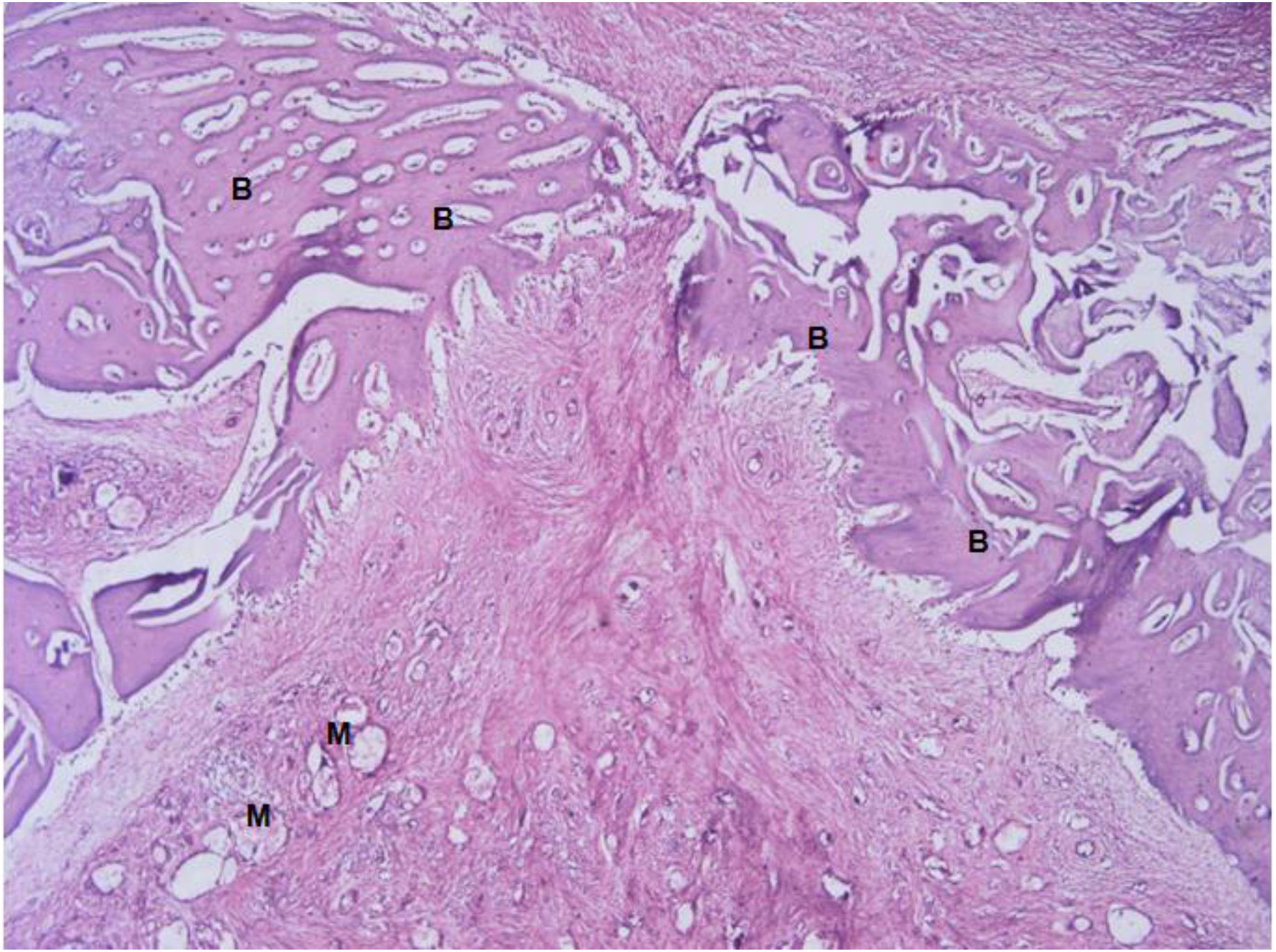
H&E stained view of the bony defect containing OsteoStat at x40 power magnification. Formation of new bone (B) can be seen on the lateral and inferior borders of the image. Haversian/volkman-like vessels are present in the central portion of the image. Fibrous stroma (F) is also present in the middle of the section at the interface of the newly formed bone.

**Fig. 2.**
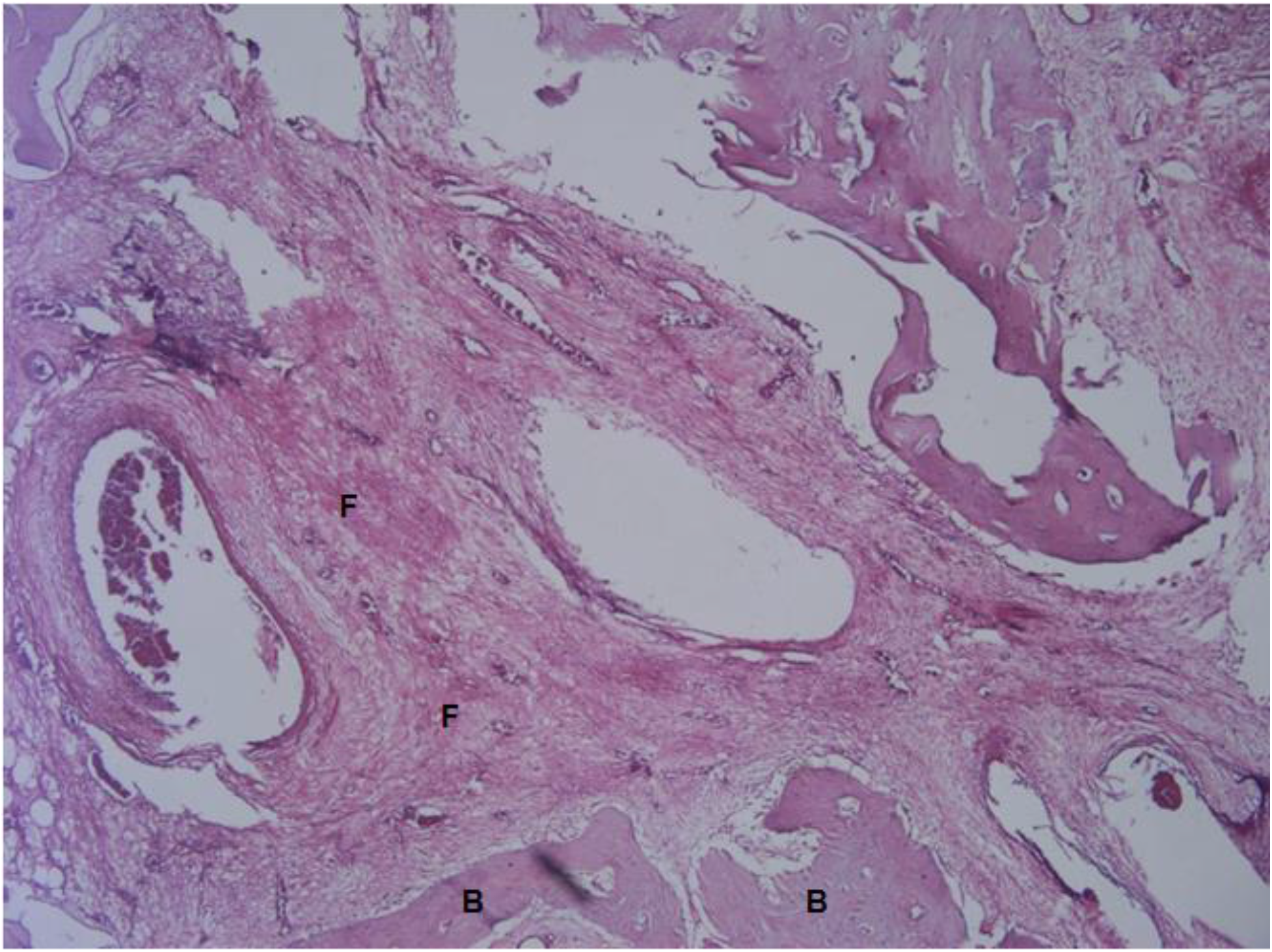
H&E section containing Bone wax at x40 power magnification. This section is dominated by fibrous stroma (F) in the middle of the image, and a small amount of new bone (B) formation at the peripheries.

#### X100 power magnification

In the OsteoStat sites, new bone formation can be seen which make up a large portions of the images **(Fig. 3).** Bone wax sites show a large amount of fibrous tissue and some new bone is also present, but as immature bony trabeculae **(Fig. 4).**

**Fig. 3.**
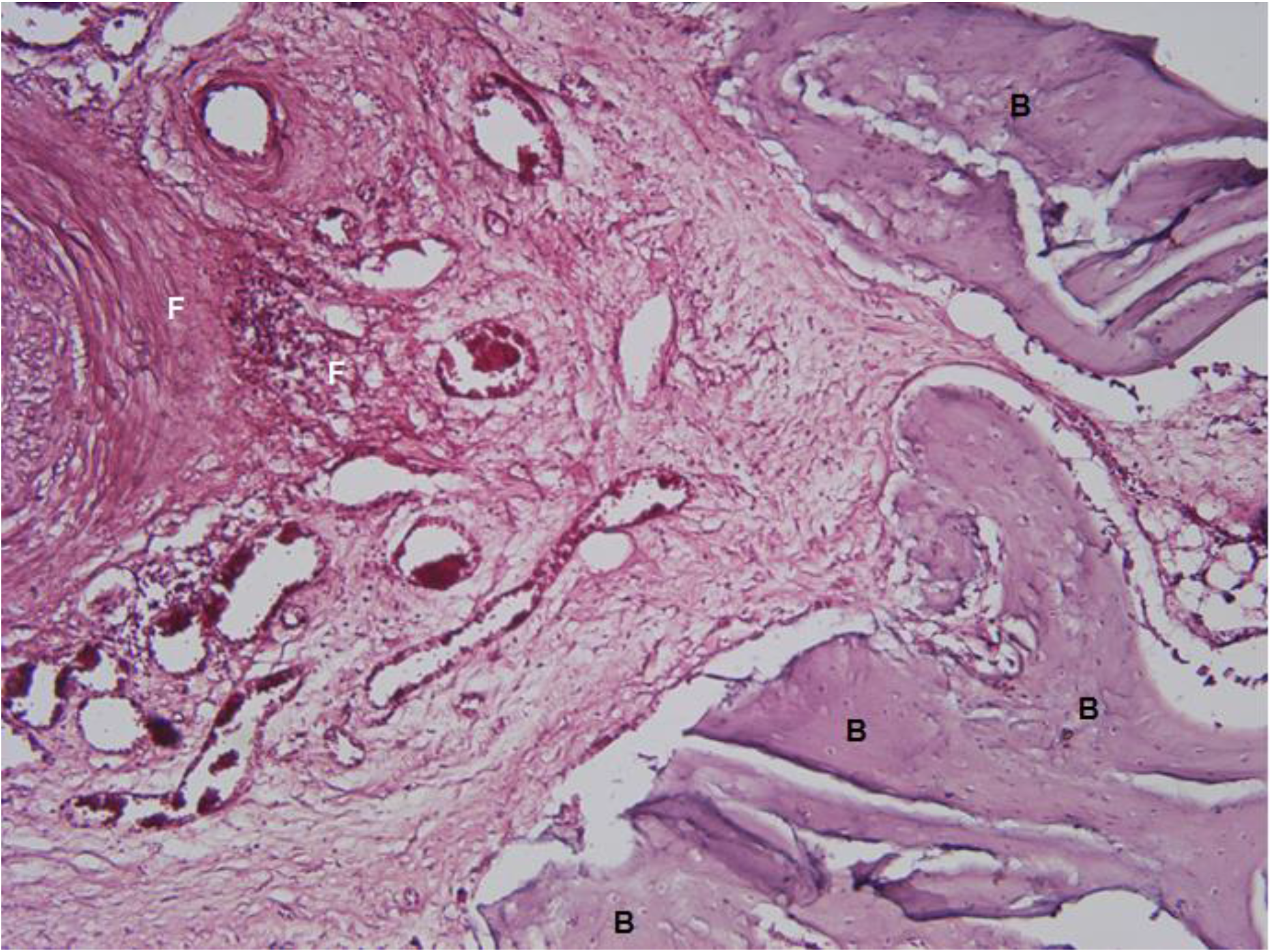
Section of OsteoStat site at x100 power magnification. New bone (B) can be seen on the right margins, with the left margin composed of fibrous stroma (F).

**Fig. 4.**
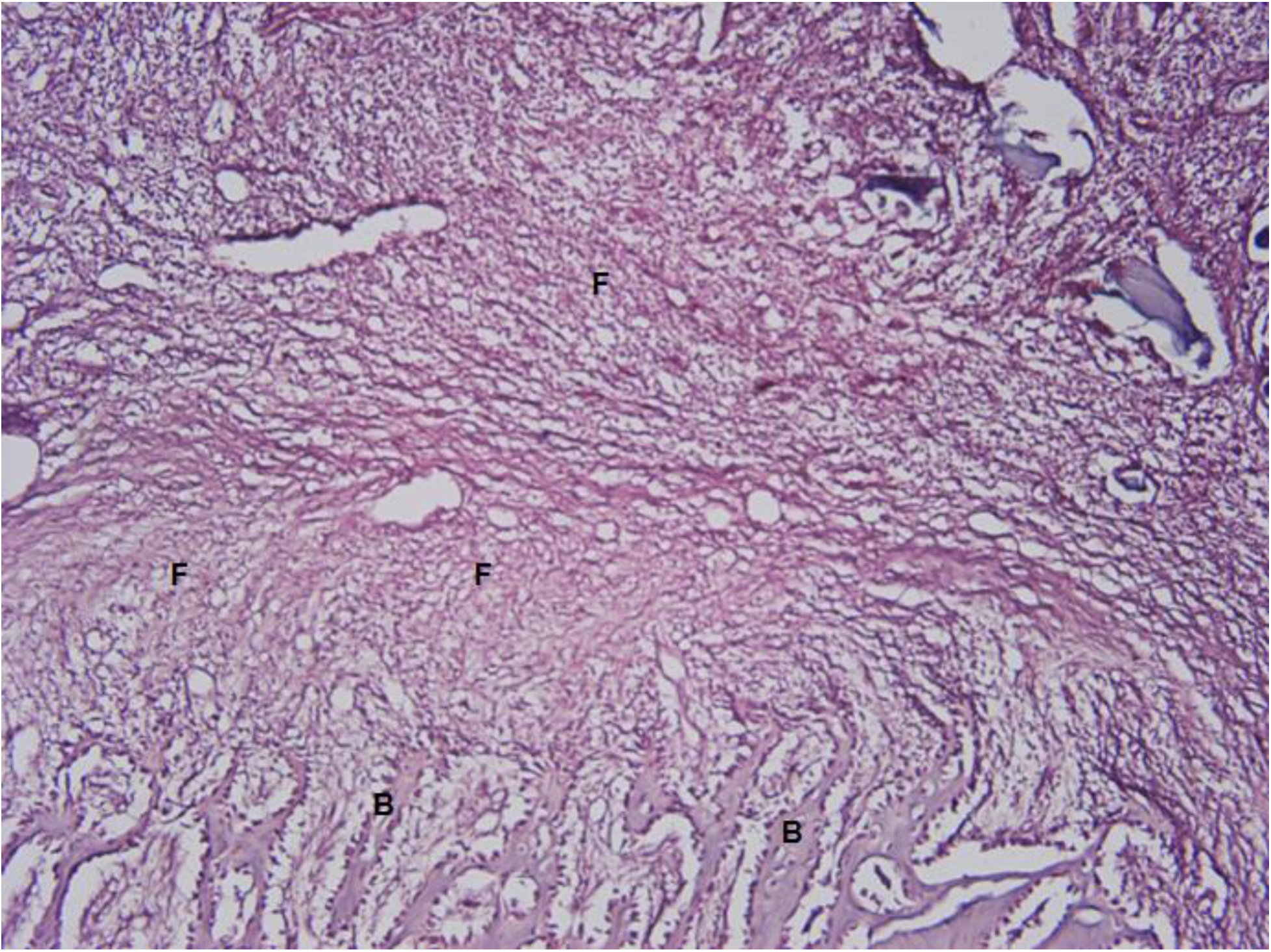
Section of the Bone wax site at x100 power magnification. Immature bony trabeculae (B) are seen at the inferior margins. The rest of the image is mainly composed of fibrous stroma (F).

#### X400 power magnification

OsteoStat sites shows active osteoblast activity lining the bony trabeculae. Osteocytes are also present in the bony matrix. No osteoclasts are seen **(Fig. 5).** Similarly, the Bone wax site **(Fig. 6)** shows osteoblast activity lining the bony trabeculae, with no osteoclasts. The variation in sections at the x400 magnification is inherently greater.

**Fig. 5.**
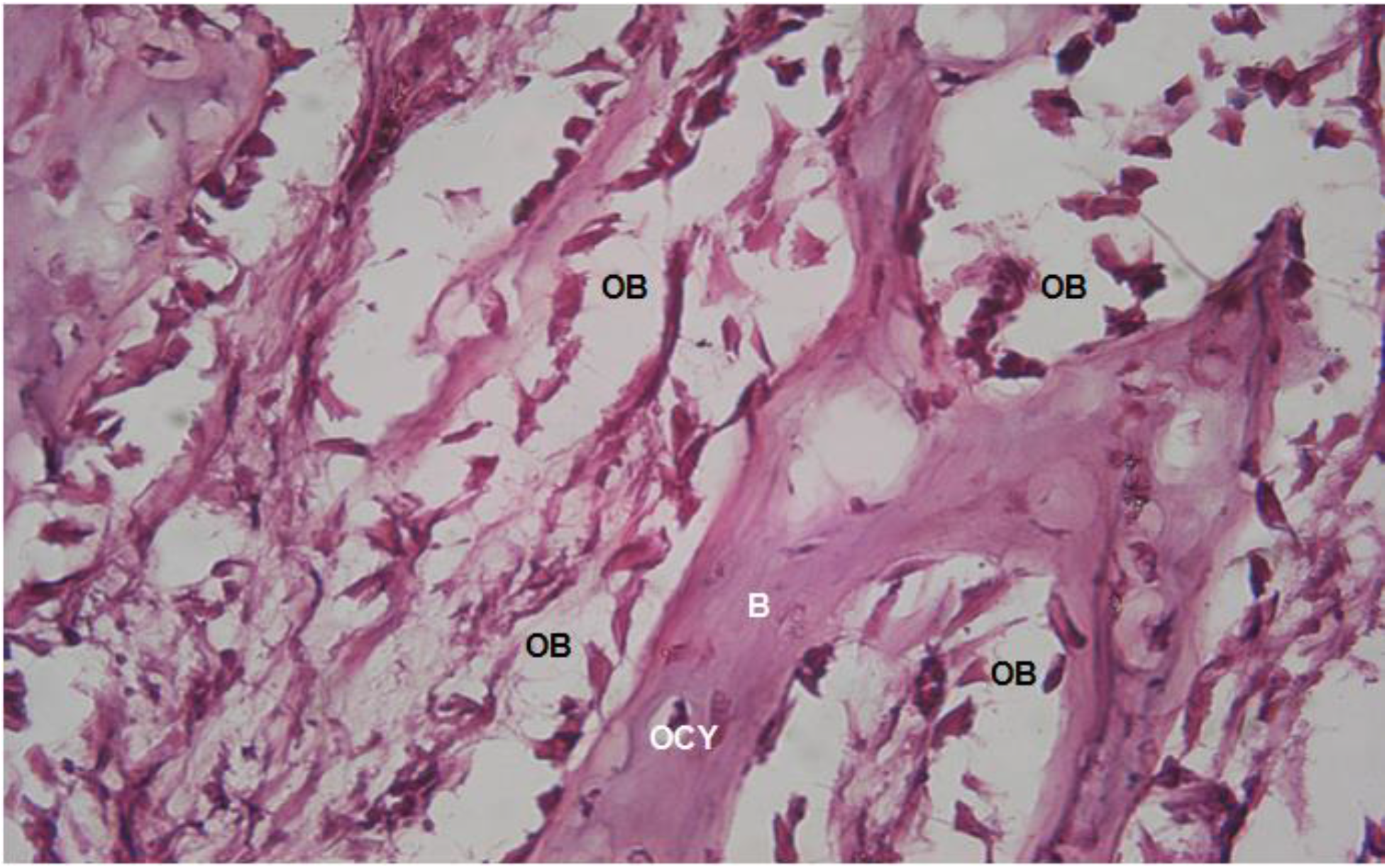
OsteoStat site at x400 power magnification. Osteoblasts (OB) are seen lining the newly formed bony trabeculae (B), with osteocytes (OCY) also present in the bony matrix.

**Fig. 6.**
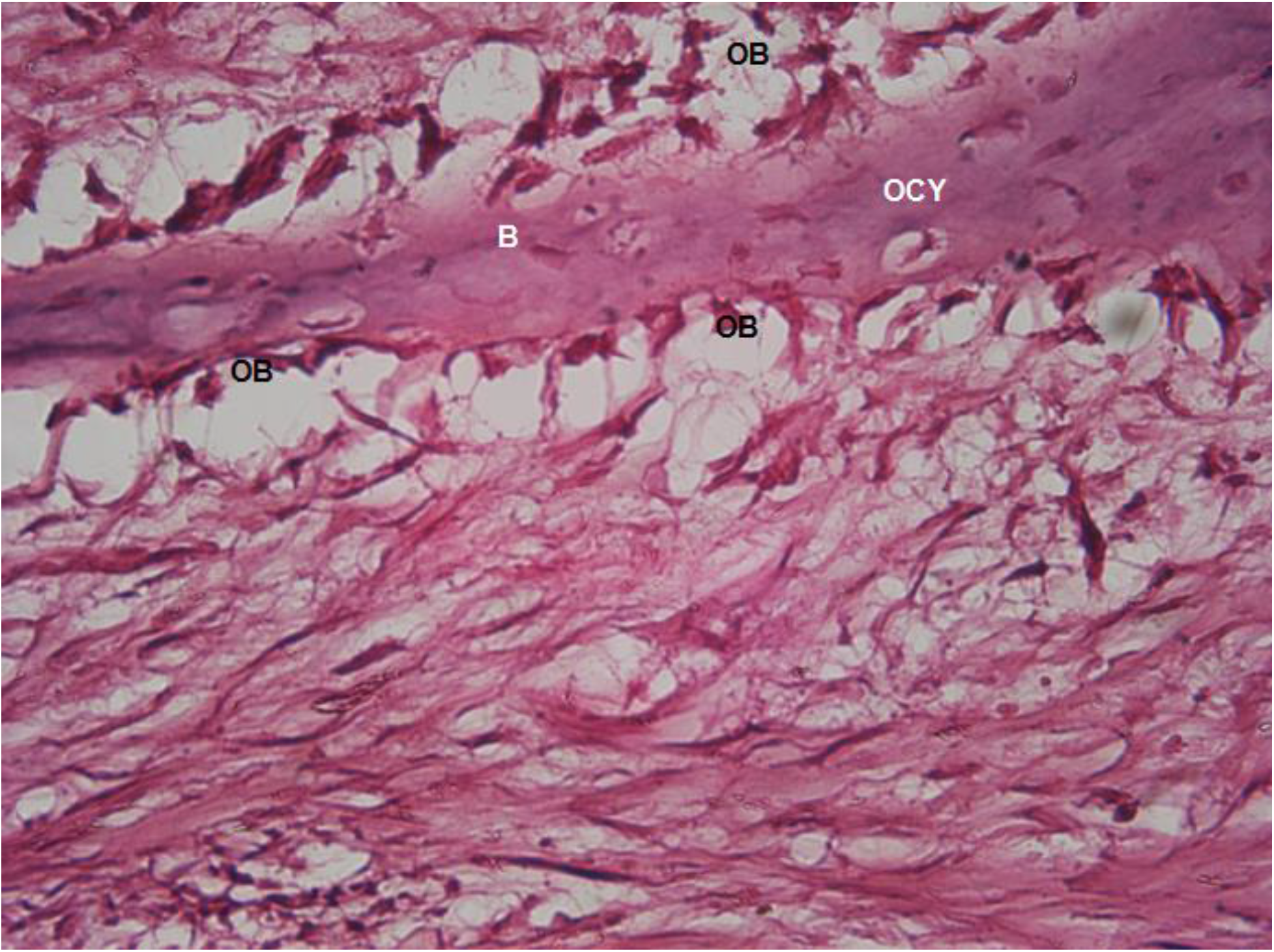
Bone wax site at x400 power magnification. Similar to figure 5, osteoblasts (OB) are seen lining the newly formed bony trabeculae (B), with osteocytes (OCY) also present in the bony matrix. The bottom portion of the image consists of soft tissue

### Histomorphometry

Histomorphometric analysis of the sections at x40 power magnification was performed. Parameters measured were mean volume fraction of test material remaining, soft tissue area and bone area. The mean area fractions of test material, soft tissue and bone in the OsteoStat composite were 12.31%, 49.64%, and 38.05% respectively. In the Bone wax group, the mean area fractions of test material, soft tissue and bone were 16.08%, 72.04%, 11.88% and respectively **(Table 1).** The difference in bone area fraction between the OsteoStat group (38.05%) and the Bone wax group (11.88%) was significant (p = 0.028) **(Table 2).** The soft tissue area fraction in the OsteoStat group (49.64%) was also less than the Bone wax group (72.04%); however, this result was not statistically significant (p = 0.089). Similarly, the amount of test material remaining in the OsteoStat group (12.31%) was less than the Bone wax group (16.08%) and was also not statistically significant (p = 0.421).

**Table 1.**
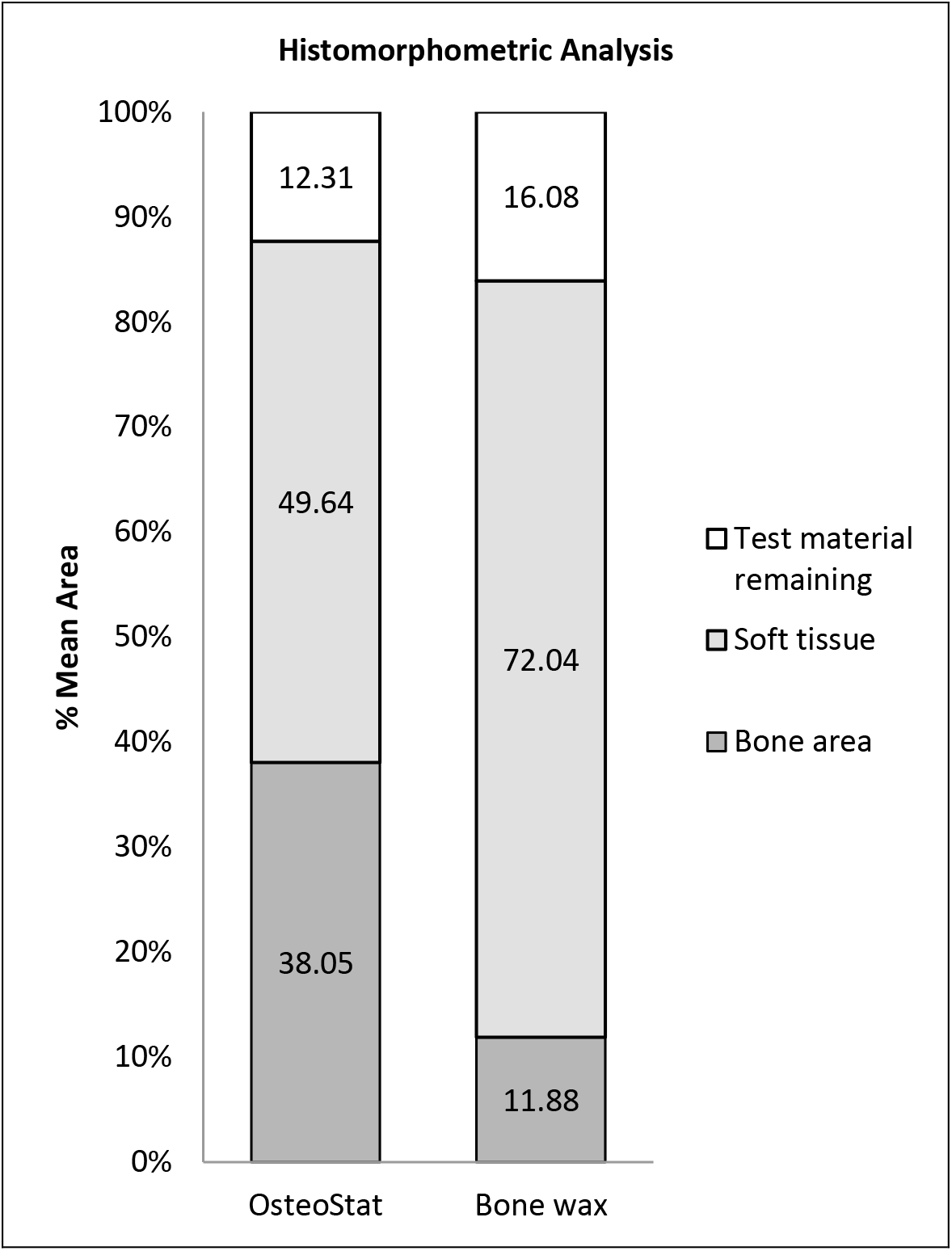
Histomorphometric Analysis of OsteoStat versus Bone wax. Histomorphometric analysis of the sections at x40 power magnification. Parameters measured were mean volume fraction of test material remaining, soft tissue and bone. All area fractions add up to make 100% of the total volume. The difference in areas of bone (p=0.028) between the two groups are statistically significant. The difference in areas of soft tissue (p=0.089) and test material remaining (p=0.421) were not statistically significant.

**Table 2.** Bone area seen in histomorphometric analysis in OsteoStat versus Bone wax. Histomorphometric analysis of 4 samples of bone defects for OsteoStat and Bone wax. The mean amount of bone observed in histomorphometric analysis was significantly greater in OsteoStat compared to bone wax. * p<0.05

| OsteoStat bone area (%) |  | Bone wax bone area (%) |  |
| --- | --- | --- | --- |
| Sample 1 | 52.49 | Sample 1 | 9.65 |
| Sample 2 | 12.42 | Sample 2 | 15.96 |
| Sample 3 | 44.31 | Sample 3 | 6.45 |
| Sample 4 | 42.98 | Sample 4 | 15.44 |
| Mean | *38.05 | Mean | *11.88 |
| Standard deviation | 17.6 | Standard deviation | 4.61 |

## Discussion

Histomorphometric data revealed that the amount of new bone was significantly greater at 6 weeks in the OsteoStat group (38.05%) versus the Bone wax group (11.88%), p=0.028. This data is consistent with the published literature in that Bone wax is a potent inhibitor of bone healing ^4,15^. Histological examinations in animal models as well as human autopsies demonstrate that the application of Bone wax not only prevents bone healing, but also promotes granuloma formation, chronic inflammation, and fibrotic scar tissue.^3,5,16^ Furthermore, Bone wax is resistant to degradation and remains in the implanted sites indefinitely.^5,16^ In a case series of 18 post-mortem examinations, histologically verified Bone wax granulomas were found, in one case as long as 10 years after implantation.^5^ Because of these potential complications, good surgical practices minimize the amount of Bone wax used, whenever needed. Its use is avoided altogether when fusion of bone is critical for post-operative function, for example, in most orthopedic surgery procedures.

Bone wax hemostatic activity is purely mechanical. It physically occludes the bleeding haversian canals in cortical and medullary bone and activates coagulation via the stasis component of Virchow’s Triad. The OsteoStat composite used in this experiment has similar handling characteristics to Bone wax, and was designed to promote hemostasis via a similar tamponade-like mechanism. Despite that no objective measures of hemostasis were designed as part of this study, both materials are subjectively reported to have equal hemostatic efficacy.

Our results showed that the amount of fibrous tissue remaining in the OsteoStat group (49.64%) was less than in the Bone wax group (72.04%), though this was not statistically significant (p=0.089). Furthermore, amount of test material remaining for the OsteoStat group (12.31%), though less than Bone wax (16.08%), was also not significant (p=0.421). One possible explanation for these findings could be that the timeframe of 6 weeks was too short for all the HA particles to take part in resorption and osteointegration. As described earlier, it would be reasonable to assume that the Bone wax particles would be resistant to degradation and resorption. However, HA particles have been shown to have complete osseous integration over a period of time, especially when blended with PLA oligomer.^17^

Subjectively, both OsteoStat and Bone wax had effective hemostatic action on the bleeding bone. However, in this experiment, we noted that OsteoStat has the added advantage of having a higher bone healing capacity compared to Bone wax. One other hemostatic Bone wax substitute that does not inhibit healing is water soluble Bone wax (WSW). WSW was first reported as

‘pluronic-based’ wax by Wang et al in 2001, and similar to this study, demonstrated superior bone healing to Bone wax.^18^ A subsequent study by Vestergaard et al of a randomized trial in humans comparing WSW versus Bone wax found no differences in infection rates, but radiologic bone measurements indicated lower levels of bone healing in Bone wax. Additionally, the study surgeons commented the WSW had some drawbacks, specifically the need to reapply due to dissolution of the WSW and the need to heat the WSW product before application to make it more pliable to smear on trabecular bone surfaces.^19^

Whether or not the lower levels of radiologic bone healing in Bone wax compared to WSW translate into real clinical effects such as sternal bone strength is a matter for debate, as demonstrated in long-term animal trials. A study comparing WSW and Bone wax in porcine sternotomies after a period of 6 months showed that although Bone wax had poorer histological and radiological outcomes, bone mechanical properties were similar. Sternal wounds closed with Bone wax were found to be weaker compared to a negative control, but no difference in sternal strength was observed between Bone wax and WSW.^20^

Similar to WSW, OsteoStat has several advantages over Bone wax and other reported Bone wax substitutes. OsteoStat used in this study has demonstrated a higher amount of bone growth, is easily sterilizable, is biocompatible and would theoretically fully integrate in bony architecture over a longer period of time.^21^ Furthermore, since it has no biological components, the material would be immunogenic and allergen free. Physical handling characteristics which are similar to Bone wax would make OsteoStat easy to use for surgeons familiar with working with traditional Bone wax.

Further studies are needed to ascertain the efficacy of OsteoStat in the human sternotomy and/or cranial/spine surgery models, particularly comparing it against Bone wax and other common hemostatic agents. Other useful avenues for investigation would be longer-term studies to ascertain the rate of HA integration in bone after application as a hemostat.

## Conclusion

This experiment has demonstrated that at 6 weeks, porcine bone defects will have higher amounts of new bone if filled with OsteoStat than with Bone wax. OsteoStat test sites also demonstrated less soft tissue and test material remaining than the Bone Wax, though the results for these parameters did not meet the threshold for statistical significance. This might be because the time frame for eventual HA osseous integration lasts many months to years. Subjectively, both OsteoStat and Bone wax have effective hemostatic properties. It is important that bone hemostat substances or its substitutes have biocompatible, osteoconductive, as well as hemostatic properties.

## List of abbreviations

PEG – polyethylene glycol, HA –hydroxyapatite, PLA-polylactic acid, mm – millimeter, H&E – hematoxylin & eosin,

## Acknowledgements

We would like to acknowledge the assistance and expertise of Liao WenJun, Wang Wei, Bao WenTao and Cheng JingWen for the animal surgery and histology preparation. We would also like to acknowledge the expertise of Molly Speltz, DVM, in the histomorphometric analysis.

## Author Contributions

Ideas; formulation or evolution of overarching research goals and aims – TT; Development or design of methodology; creation of models – TT; Verification, whether as a part of the activity or separate, of the overall replication/reproducibility of results/experiments and other research outputs - TT PC JS JB KR; Application of statistical, mathematical, computational, or other formal techniques to analyse or synthesize study data – TT; Conducting a research and investigation process, specifically performing the experiments, or data/evidence collection – TT; Provision of study materials, reagents, materials, patients, laboratory samples, animals, instrumentation, computing resources, or other analysis tools - TT PC JS JB KR; Management activities to annotate (produce metadata), scrub data and maintain research data (including software code, where it is necessary for interpreting the data itself) for initial use and later re-use – TT; Preparation, creation and/or presentation of the published work, specifically writing the initial draft (including substantive translation) – TT; Preparation, creation and/or presentation of the published work by those from the original research group, specifically critical review, commentary or revision – including pre- or post-publication stages – TT; Preparation, creation and/or presentation of the published work, specifically visualization/data presentation - TT PC JS JB KR; Acquisition of the financial support for the project leading to this publication - TT PC JS JB KR

